# Endosperm and seed transcriptomes reveal possible roles for small RNA pathways in wild tomato hybrid seed failure

**DOI:** 10.1101/2019.12.20.884387

**Authors:** Ana Marcela Florez-Rueda, Flurin Fiscalini, Morgane Roth, Ueli Grossniklaus, Thomas Städler

**Author notes:** Author for Correspondence: Ana Marcela Florez-Rueda, Department of Plant and Microbial Biology & Zurich–Basel Plant Science Center, University of Zurich, 8008 Zurich, Switzerland.

## Abstract

Crosses between the wild tomato species *Solanum peruvianum* and *S. chilense* result in hybrid seed failure (HSF), characterized by endosperm misdevelopment and embryo arrest. We previously showed that genomic imprinting, the parent-of-origin–dependent expression of alleles, is perturbed in hybrid endosperm, with many of the normally paternally expressed genes losing their imprinted status. Here, we report transcriptome-based analyses of gene and small RNA expression levels. We identified 2,295 genes and 468 small RNAs (sRNAs) as differentially expressed (DE) when comparing reciprocal hybrid seed to seeds and endosperms from the two within-species crosses. Our analyses uncovered a pattern of overdominance in endosperm gene expression in both cross directions, in marked contrast to the patterns of sRNA expression in whole seeds. Intriguingly, patterns of increased gene expression resembled the previously reported increased maternal expression proportions in hybrid endosperms. We identified physical clusters of sRNAs; DE sRNAs exhibited reduced levels of expression in hybrid seeds from both cross directions. Moreover, sRNAs mapped to genes coding for key proteins involved in epigenetic regulation of gene expression, suggesting a regulatory feedback mechanism. We describe examples of genes that are targets of sRNA-mediated gene silencing; in these cases, reduced sRNA expression was concomitant with increased gene expression in hybrid seeds. Our analyses also show that *S. peruvianum* dominance impacts gene and sRNA expression in hybrid seeds. Overall, our study indicates roles for sRNA-mediated epigenetic regulation in HSF between closely related wild tomato species.

## Introduction

The establishment of reproductive barriers between diverging lineages is a basic component of the speciation process and thus of major interest in evolutionary biology (Coyne and Orr 2004). In this study, we assess the molecular correlates of hybrid seed failure (HSF), a form of postzygotic barrier acting early in seed development of many flowering plants. In the angiosperm seed, embryo and endosperm are the products of two independent fertilization events. The endosperm is usually a triploid tissue that nourishes the growing embryo; failure of proper endosperm development often leads to embryo arrest and is considered the main cause of HSF (Rebernig et al. 2015; Garner et al. 2016; Oneal et al. 2016). HSF has been frequently observed upon hybridization of closely related homoploid plant species as well as between lineages differing in ploidy (Beamish 1955; Rick 1963; Scott et al. 1998; Dilkes et al. 2008; Jullien and Berger 2010; Lu et al. 2012; Burkart-Waco et al. 2013; Rebernig et al. 2015; Oneal et al. 2016).

From an evolutionary perspective, the developing seed can be viewed as an arena in which the two parental genomes ‘collide’. Any differences in parental optima for resource allocation to progeny (representing parental conflict) are expected to manifest in the endosperm (Haig and Westoby 1991; Haig 2013). The ratio of ‘effective’ parental genomic contributions in the endosperm appears to largely determine the success or failure of particular crosses, an interpretation bolstered by the frequent observation that postzygotic barriers can be weakened by manipulating the ploidy of one of the parents (Johnston et al. 1980; Josefsson et al. 2006; Lafon-Placette and Köhler 2016). Transgressive and complementary hybrid seed phenotypes are common and thought to reveal different levels of parental conflict between lineages (Lu et al. 2012; Haig 2013; Rebernig et al. 2015; Florez-Rueda, Paris, et al. 2016). These observations have led to the hypothesis that parent-of-origin–dependent allelic expression (i.e. genomic imprinting) might be causally involved in HSF. Genomic imprinting is an epigenetic phenomenon causing the preferential expression of alleles depending on their parental origin. In flowering plants, while occurring also in the embryo, genomic imprinting is prevalent in the endosperm and critical for proper seed development (Berger 2003).

Although perturbed genomic imprinting has been shown to be a molecular correlate of HSF (Gutierrez-Marcos et al. 2003; Josefsson et al. 2006; Walia et al. 2009; Jullien and Berger 2010; Wolff et al. 2015; Florez-Rueda, Paris, et al. 2016; Lafon-Placette et al. 2018), successful seed development results from the precise orchestration of additional genomic and developmental processes. Other molecular processes during seed formation, like de-repression of transposable elements (TEs; Fultz et al. 2015; Martínez and Köhler 2017) and gene regulation mediated by small RNAs (sRNAs; Bourc’his and Voinnet 2010; Ng et al. 2012; Lu et al. 2012) likely act in the endosperm to determine the success or failure of particular cross combinations. Of particular interest are sRNAs; these RNA forms are involved in plant development, reproduction, and genome reprogramming (Haig 2013; Benkovics and Timmermans 2014; Borges and Martienssen 2015; Martínez and Köhler 2017).

For instance, microRNAs (miRNAs) are post-transcriptional regulators of gene expression, and various other types of sRNAs are involved in post-transcriptional gene silencing (PTGS) via transcript cleavage or translational repression as well as in transcriptional gene silencing (TGS), the latter mostly via RNA-directed DNA methylation (RdDM; Matzke and Mosher 2014; Pikaard and Mittelsten Scheid 2014; Borges and Martienssen 2015; Cuerda-Gil and Slotkin 2016; D’Ario et al. 2017). Several recent studies point to a pivotal role for sRNA-mediated gene silencing in regulating proper seed development and/or hybrid fitness (Groszmann et al. 2011; Lu et al. 2012; Rodrigues et al. 2013; Vu et al. 2013; Martínez et al. 2016, 2018; Borges et al. 2018). Although our current knowledge regarding sRNA biogenesis and regulatory mechanisms stem mainly from work in the model species *Arabidopsis thaliana*, it is expected that the underlying concepts apply to most angiosperms, albeit some deviations from the canonical mechanisms may occur in more distantly related taxa, such as our model system *Solanum*.

In this study, we quantified the expression patterns of sRNAs in reciprocal crosses between two wild tomato species that show near-complete HSF, an important postzygotic barrier to interbreeding among several species of wild tomatoes (*Solanum* section *Lycopersicon*). Classical studies found high proportions of HSF in reciprocal crosses between the closely related *S. peruvianum* (P) and *S. chilense (C)* (Rick and Lamm 1955). Following this pioneering work, we have quantified various degrees of seed inviability in reciprocal hybrid crosses involving several species of wild tomatoes. Moreover, we observed differences in the cellular architecture and histology of failing endosperms, as well as strong differences in seed size depending on the direction of hybrid crosses (Roth, Florez-Rueda, Griesser, et al. 2018). Similar HSF-associated phenotypes have been described in different *Solanum* species and other angiosperm taxa, including interploidy and homoploid hybrid crosses in model species and important crops (Cooper and Brink 1945; Beamish 1955; Rick 1963; Ortiz and Ehlenfeldt 1992; Scott et al. 1998; Dilkes et al. 2008; Jullien and Berger 2010; Ishikawa et al. 2011; Burkart-Waco et al. 2013; Rebernig et al. 2015).

We previously studied the molecular correlates of HSF in the *S. peruvianum*–*S. chilense* case and found that genomic imprinting in the endosperm is systematically perturbed (Florez-Rueda, Paris, et al. 2016), but we did not assess changes in overall expression levels. This intriguing pattern motivated us to investigate the likely epigenetic basis of strong HSF as observed in *S. peruvianum*–*S. chilense* crosses, with a focus on the possible roles of sRNAs. In the present study, we integrate gene and sRNA expression estimates and study their expression profiles in both normally developing and failing hybrid endosperm and seeds, respectively. We examine the sRNAs’ targets and provide examples of representative genes exhibiting changes in gene expression concomitant with sRNA expression variation. By comparing the expression of hybrids and their parents, we further test how expression inheritance patterns are shaped by different ‘effective ploidies’ of the parental lineages.

## Materials and Methods

### Plant Material, RNA Extraction, and Library Preparation

All seeds were obtained from the C.M. Rick Tomato Genetics Resource Center at U.C. Davis (http://tgrc.ucdavis.edu, last accessed 16 June 2016). For *S. peruvianum*, we used seeds from accession LA1616 (Dept. Lima, Peru) and for *S. chilense*, we used seeds from accession LA4329 (Region Antofagasta, Chile). We used four plants, referred to as 1616A, 1616J, 4329B, and 4329K, and analyzed three different parental combinations: the within-species *S. peruvianum* case (PP) with plants 1616A and 1616J as parents, the within-species *S. chilense* (CC) case with plants 4329B and 4329K as parents, and the hybrid cases (PC and CP) with plants 1616A and 4329B in both parental roles in reciprocal crosses. The parental plants were grown from seeds and transferred to a climate chamber before the onset of the experiments. The conditions in the climate chamber were 12 h light (18 klux) at 22°C with 50% relative humidity and 12 h darkness (0 klux) at 18°C with 60% relative humidity. For each of the three cross types, hand pollinations were performed and developing fruits were collected on each plant for each cross type.

Based on prior seed development studies in *Solanum* (e.g. Beamish 1955; Briggs 1993) and our own histological analyses (Roth, Florez-Rueda, Griesser, et al. 2018), we chose an early globular embryo stage to collect the material for library preparation. We collected fruits 14 days after pollination (DAP), always in the late afternoon. This developmental stage was chosen because it was early enough to distinguish the developing embryo from the surrounding endosperm tissue, while the latter was large enough to extract RNA in the quantities needed for library preparation. For each plant and cross type, two separate RNA libraries were prepared from endosperm tissue, for a total of 12 endosperm libraries. The raw data for the endosperm transcriptomes has been published before, and detailed methodology for its production was described in Florez-Rueda, Paris, et al. (2016). In brief, fruits were harvested, fixed, and endosperms were laser captured with the Laser-Assisted Microdissection (LAM) technique outlined in Florez-Rueda, Grossniklaus, et al. (2016).

The same crossing design described above for endosperm transcriptomes was implemented for the whole-seed sRNA data sets. As we were interested in overall—rather than parent-specific—sRNA expression levels and sRNAs were found to be abundant in all three *Arabidopsis* seed compartments (Erdmann et al. 2017; Kirkbride et al. 2019), we extracted sRNAs from whole seeds. Moreover, we generated sRNA libraries only from hybrid and normal seeds from plants 1616A and 4329B, serving as parents in both intra- and inter-specific crosses (supplementary figure S1, Supplementary Material online). For these sRNA libraries, we used three replicates for our analyses, each replicate reflecting independent hand-pollination events performed on different days. As for the endosperm transcriptomes, developing fruits were collected at 14 DAP in the late afternoon and immediately placed into RNAlater solution. The samples were immediately transferred to a refrigerator and remained in the RNAlater solution for a minimum of 24 h and a maximum of 48 h. Whole seeds were dissected in RNase-free water and subjected to consecutive water washes to remove the fruit flesh debris. We collected a minimum of 1 µg of seeds from tens of fruits from each cross type and proceeded to sRNA extraction. RNA was extracted using the miRVana RNA isolation kit (Ambion, Life Technologies Corporation, Foster City, CA, USA). sRNA libraries were prepared using the NEXTflex SRNAs-Seq Kit v2 according to the manufacturer’s protocol (Bioo Scientific Corporation, Austin, TX, USA). Libraries were sequenced in single-end fashion on one lane of an Illumina HiSeq 4000 at the Functional Genomics Center Zurich (www.fgcz.ch).

### Read Mapping and Differential Expression Analyses

Mapping of sRNA reads was performed using STAR (Dobin et al. 2013) with a maximum of two mismatches to the SL2.50 assembly of the cultivated tomato reference genome (The Tomato Genome Consortium 2012) deposited in ensemble genomes (https://plants.ensembl.org/Solanum_lycopersicum/Info/Annotation/#genebuild, last accessed 7 February, 2017). After using Shortstack (Axtell 2013), 122,398 sRNA clusters were identified with default options. Of these, 57,711 fell within the coordinates of a gene coding region or its 2.5 kb flanking regions using BEDTools window command (Quinlan and Hall 2010); these clusters were used for further analyses. Of these 57,711 clusters and based on the corresponding SL2.50 ensemble annotation of the genome, we annotated 7,112 sRNAs as miRNAs; Shortstack inferred 44 additional miRNAs for a total number of 7,156 miRNAs. Other forms of non-coding RNAs that were represented in our sRNA libraries included 2,949 antisense RNAs, rRNAs, tRNAs, snoRNAs, snRNAs, and SRPRNAs. These latter forms were removed before performing differential expression analyses. By using the counts provided by Shortstack, we performed differential gene expression analyses using DESeq2 (Love et al. 2014) in the same manner as for the endosperm transcriptomes (see below).

We reanalyzed the endosperm transcriptome data previously produced (Florez-Rueda, Paris, et al. 2016). Raw reads were mapped to the SL2.50 assembly of the tomato genome deposited in ensemble genomes (https://plants.ensembl.org/Solanum_lycopersicum/Info/Annotation/#genebuild). The tuxedo pipeline (Trapnell et al. 2012) was used for the assembly of reads, mapping to the tomato reference genome, and count estimation. Raw count tables were produced with additional packages of the Tuxedo pipeline, cuffquant and cuffnorm; unnormalized counts per transcript were used for subsequent analyses. Differential gene expression analyses for transcripts as well as for sRNA clusters were performed using DESeq2 (Love et al. 2014), as implemented in the RNAseqWrapper package (Schmid 2017) in R (R Development Core Team 2014). To test for differential gene expression between viable and hybrid seeds while taking into account expression variation within both species, a model of a single factor with multiple levels (species correspondence: *S. peruvianum*, *S. chilense*, and type of seed: normal, hybrid) was implemented in the given RNAseqWrapper module (Schmid 2017). This implies that we contrasted all within-species expression data as one entity (from crosses PP, CC) with all hybrid expression data as the other entity (from crosses PC, CP). However, to identify possible differences between species, we additionally considered separate within-species comparisons for the DE sRNA clusters (i.e. PP *vs*. PC, and CC *vs*. CP). While our main focus is the comparison between intraspecific and failing hybrid seeds, we thus additionally report all DE sRNAs from the two separate contrasts. DE transcripts and sRNAs with more than absolute 2.5 log fold-change and a Bonferroni-corrected *P* value <0.05 are reported as significantly DE.

Downstream gene enrichment analyses were carried out using the STRING database (Szklarczyk et al. 2017). We report functional enrichment analyses from STRING with a False Discovery Rate (FDR) of 0.01. When reported, GO assignment was assessed using the PANTHER database (Mi et al. 2017). These two databases, STRING and PANTHER, were also used for fine-tuning annotation of genes that lacked annotation in the corresponding SL2.50 ensemble functional annotation files. In the set of DE sRNAs, we defined clusters of genes along the chromosomes based on their curated joint annotation; they were defined as a gene cluster if three or more genes with the same annotation were located within 5 Kb of genomic space.

### Expression Mode Classification

We compared endosperm expression levels of transcripts and sRNA clusters among *S. peruvianum* (PP), *S. chilense* (CC), and their reciprocal hybrids (PC and CP), following the rationale of previous studies to discriminate among the various categories of expression modes (McManus et al. 2010; Combes et al. 2015). Independent of whether a gene or sRNA was found to be DE, genes with less than one-fold change between hybrid and normal endosperm were considered as showing *conserved* expression; for sRNAs, we used a lower threshold of 0.5-fold expression change. The mode of expression was determined as *additive* if the expression in the hybrid was less than in *S. peruvianum* and greater than in *S. chilense* (or vice versa). If the expression in the hybrid endosperm was similar as in one of the parental species it was classified as *dominant* for the respective species, and genes and sRNAs with either higher or lower hybrid expression than in both *S. peruvianum* and *S. chilense* were classified as exhibiting *overdominant* and *underdominant* expression, respectively.

## Results

### Mapping and Gene Identification

We performed sRNA sequencing from whole seeds obtained from intra- and reciprocal inter-specific crosses. Three replicate sets of ‘normal’ and ‘hybrid’ sRNA transcriptomes were produced for each of the two main parental plants, the same individuals that were used in our previous study (supplementary figure S1, Supplementary Material online; Florez-Rueda, Paris, et al. 2016). After sequencing, we obtained a mean of 9.6 million reads per library, of which a mean of 7.7 million reads were kept after quality filtering and mapping (supplementary table S1, Supplementary Material online). Not surprisingly, the 24-nt sRNA category was the prevalent type among the 57,711 clusters identified within 2.5 Kb of annotated genes, whereas 21-22-nt sRNAs and miRNAs accounted for only 10% of the total identified sRNA clusters (supplementary table S2, Supplementary Material online). To integrate sRNA and gene expression information, we remapped our previously produced endosperm transcriptomes obtained after LAM (Florez-Rueda, Paris, et al. 2016) to the *S. lycopersicum* reference genome. A mean of 21 million reads per library mapped uniquely to the reference genome and was used in subsequent analyses, making the mean proportion of retained reads 84% of the initially obtained raw data (supplementary table S1, Supplementary Material online). We thus detected 33,805 transcripts across all endosperm transcriptomes.

### Differential Expression in Hybrid Endosperms of Wild Tomatoes

We identified common trends of differential expression between normal and hybrid endosperms obtained from both within-species and hybrid crosses, with LA1616A (P) and LA4329B (C) serving as maternal parents in both cross types [contrast (PP, CC) *vs.* (PC, CP)]. Genes that are consistently DE in the hybrid endosperms of both species tend to have higher levels of expression when compared to ‘normal’ (intraspecific) endosperms in each species (figure 1A). Of the 33,805 transcripts for which we obtained expression values, 2,295 were found as DE in hybrid endosperms; transcripts identified as DE are reported in supplementary table S3, Supplementary Material online. Of these, 1,515 were found overexpressed and 780 underexpressed in hybrid compared to normal endosperms from the same maternal plants.

**FIG. 1.**
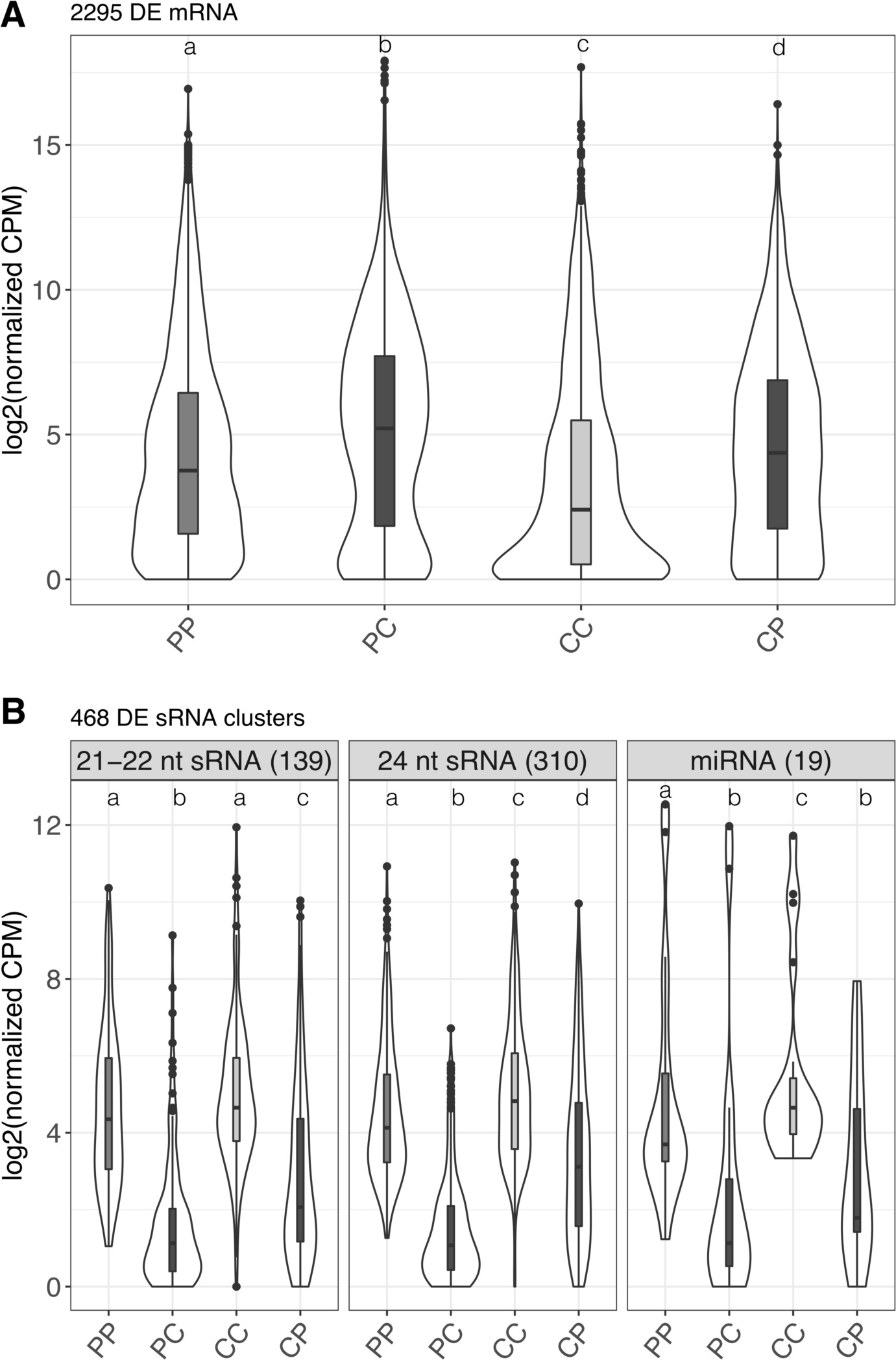
Expression distributions of 2,295 differentially expressed genes in the endosperm (*A*), and 468 sRNA clusters in whole seeds (*B*) differentially expressed between within-species and hybrid crosses [contrast (PP, CC) *vs.* (PC, CP)]. Plants LA1616A (P) and LA4329B (C) served as maternal plants in both cross types. Letters on top of violin plots represent significant differences between expression distributions (Wilcoxon rank-sum test, *P* < 0.01).

To test possible roles of sRNAs in mediating HSF and the increases in gene expression (figure 1A) and maternal allelic proportions (Florez-Rueda, Paris, et al. 2016), we investigated patterns of sRNA expression. The pattern of whole-seed sRNA differential expression is in stark contrast to the increase in gene expression we found among DE genes in hybrid endosperms. From the 57,229 total sRNA clusters identified across all sRNA libraries, 468 clusters were DE. These corresponded to miRNAs (19), 24-nt sRNAs (310), and 21-22-nt sRNAs (139) (figure 1B; supplementary table S4, Supplementary Material online). Their altered expression is consistent in reciprocal hybrid crosses, with sRNAs being underexpressed in both PC and CP hybrid seeds. The magnitude of the differences in sRNA expression is larger in seeds from *S. peruvianum* maternal plants (figure 1B), thus mirroring the differences in seed phenotype and increases in maternal allelic proportions in hybrid endosperms, which are both more marked in hybrid seeds with *S. peruvianum* as the seed parent (Florez-Rueda, Paris, et al. 2016; Roth, Florez-Rueda, Griesser, et al. 2018).

To shed light on the roles of a putative Pol lV sRNA pathway in *Solanum* (see Erdmann et al. 2017), we examined patterns of expression of the principal subunits of Pol IV, Pol V, and Pol II in the endosperm of hybrid *vs* normally developing *Solanum* seeds (supplementary table S5, Supplementary Material online). We observed reduced expression in hybrids of both genes coding for the subunits of Pol lV: RNA polymerase 4 second largest subunit, *RPD2* (logFC = –0.56, FDR-corrected *P* = 0.0148) and RNA polymerase 4 largest subunit, *RPD1* (logFC = –1.82, FDR-corrected *P* = 9.69E-49), as well as reduced expression of the gene coding for subunit H of RNA polymerase V (logFC= – 2.09, FDR-corrected *P* = 1.75E-55).

Following the general trend of increased expression in hybrid endosperms (figure 1A), PF00067 Cytochrome P450 was enriched with 40 genes found DE (supplementary figure S2*A*, supplementary table S6, Supplementary Material online). Some of these genes are part of the brassinosteroid biosynthesis KEEG pathway, which is also enriched in the set of genes that show overexpression in hybrid endosperms. Eight genes belonging to the ethylene biosynthetic pathway (GO:0009693) consistently showed increased expression in hybrid endosperms, including *ACO3*, *ACO4*, *ACO1*, and *ACS3*, among other homologs of carboxylate synthases and oxidases in ethylene production (supplementary tables S3 and S6, Supplementary Material online). In addition to genes involved in ethylene metabolism showing increased expression, 15 ethylene-responsive TF genes were also overexpressed in hybrid endosperms (supplementary figure S2*C*, Supplementary Material online), contributing to the enrichment of the IPR00147-AP2/ERF domain (supplementary table S6, Supplementary Material online). Genes belonging to this family include highly conserved imprinted genes in *Solanum* and other plant species (Ikeda 2012; Florez-Rueda, Paris, et al. 2016; Roth, Florez-Rueda, Paris, et al. 2018).

The general pattern of overexpression in the hybrid endosperms holds particularly for TF genes (supplementary table S6, supplementary figure S2*C*–*F*, Supplementary Material online). Genes encoding subunits of the mediator complex, a global regulator of polymerase II, were found overexpressed in hybrid endosperms, with the term IPR013921 mediator complex significantly enriched. Overexpression was much higher in hybrid endosperm of *S. chilense* than of *S. peruvianum* maternal plants (supplementary figure S2*D*, Supplementary Material online), with many of these genes belonging to the term GO:0001104, RNA polymerase II transcription cofactor activity. Strikingly, we uncovered the consistent overexpression of 29 genes containing a MADS-box (IPR002100), likewise displaying more substantial increases of gene expression in hybrid seeds with *S. chilense* maternal plants (supplementary table S6, supplementary figure S2*E*, Supplementary Material online).

### Joint Signatures of Gene and sRNA Expression Dynamics

To investigate the potential role of sRNAs in modulating gene expression in the endosperm, we integrated our seed sRNA data with our endosperm transcriptome data. sRNAs were given the annotation of the gene they mapped to if they fell within 2.5 kb boundaries (supplementary table S4, Supplementary Material online). Strikingly, the identity of many genes with mapped DE sRNAs revealed roles in epigenetic regulation and/or sRNA biogenesis, suggesting a regulatory feedback mechanism. We identified 119 DE sRNA clusters targeting 79 genes in which underexpression of sRNAs in hybrid seeds was concomitant with the overexpression of the corresponding genes in hybrid endosperms of both cross directions, PC and CP (figure 2). These particular cases suggest gene silencing by the reported clusters of sRNAs that is partly defective in hybrid seeds. Two of these identified genes (Solyc02g091030, Solyc05g012640) encode proteins with RNA and DNA polymerase activity, respectively, and are highly expressed in normal tomato endosperm (figure 2*A*, *B*). The tomato homolog of *DEFECTIVE IN MERISTEM SILENCING3* (*DMS3*), a component of the canonical RdDM pathway in *Arabidopsis* (Matzke and Mosher 2014), is targeted by DE sRNA clusters in both species (figure 2C), as is a gene encoding a REMORIN protein (Solyc06g035920; figure 2D).

**FIG. 2.**
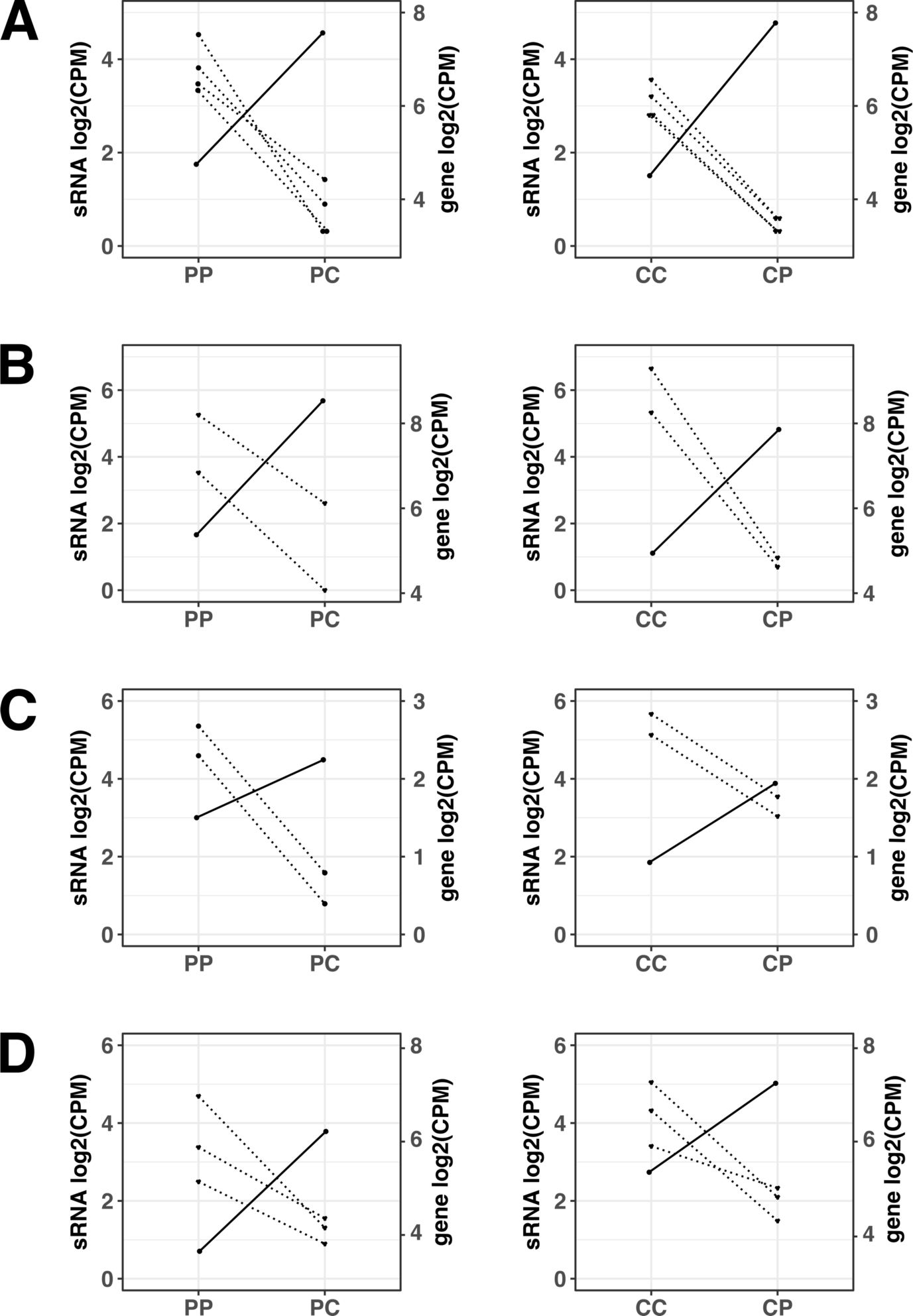
Examples of decreased sRNA expression being concomitant with gene overexpression in both reciprocal hybrids, indicating that sRNA-mediated gene silencing may be involved. (*A*) Solyc02g091030, nuclear transcription factor Y subunit C-2 with DNA-directed DNA polymerase activity. (*B*) Solyc05g012640, T7 RNA polymerase gene 1. (*C*) Solyc03g083120, *DEFECTIVE IN MERISTEM SILENCING3*, *DMS3*. (*D*) Solyc06g035920, *Remorin-1.* Dot plots show patterns of expression; on the left x axis small RNAs expression and on the right x axis gene expression. Left and right panels show normal and hybrid seeds with *S. peruvianum* (P) and *S. chilense* (C) as maternal plants, respectively. Each dot represents an sRNA cluster and lines within single dot plots trace changes in expression between normally developing and hybrid seeds. Dotted lines and solid lines trace sRNA and gene expression, respectively. All sRNA clusters are significantly DE. Not all genes shown are significantly DE but do have positive fold-changes, indicating gene overexpression in hybrid seeds.

Furthermore, we identified 75 genes targeted by 113 sRNA clusters with asymmetrical changes in gene expression concomitant with reduced sRNA expression in reciprocal PC and CP hybrids (figure 3), highlighting cases of dissimilar epigenetic responses to hybridization. The gene Solyc10g005160, *PURINE PERMEASE4* (*PUP4*) and two clustered *LATERAL ORGAN BOUNDARIES* (*LOB*, Solyc09g014700 and Solyc09g014690) genes showed strongly increased expression in the CP hybrid endosperm while their expression in PC hybrid endosperm remained low, concomitant with reduced sRNA expression (figure 3A, B). In contrast, Solyc08g007530, *AT-HOOK MOTIF NUCLEAR-LOCALIZED PROTEIN1* (*AHL1*), showed reduced gene expression in CP hybrid but increased gene expression in the PC hybrid endosperm (figure 3C), similar to patterns of gene expression of Solyc03g098280 (*SlAGO1b*), an *ARGONAUTE 1b* gene (figure 3D). ARGONAUTE proteins are core components of the sRNA-dependent silencing pathways (Matzke and Mosher 2014; Pikaard and Mittelsten Scheid 2014). Six sRNA clusters mapped within the boundaries of this gene; they were less expressed in hybrid seeds, concomitant with higher gene expression in the PC hybrid but slightly decreased gene expression in CP hybrid endosperm (figure 3D).

**FIG. 3.**
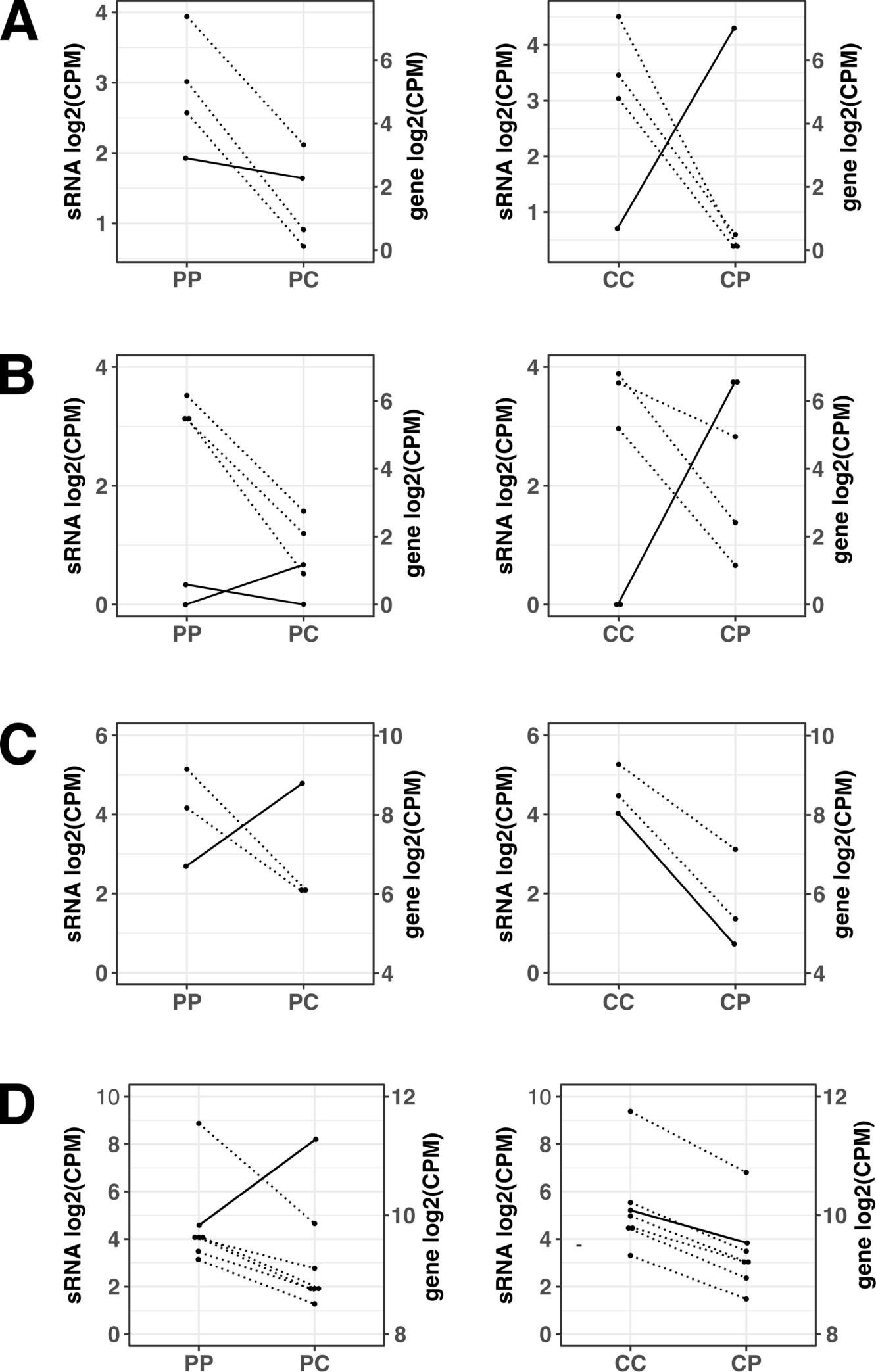
Examples of sRNA downregulation being concomitant with asymmetrical changes in gene expression in the reciprocal hybrids. (*A*) Solyc10g005160, *PUP4*. (*B*) Solyc09g014700, Solyc09g014690, *LOB.* (*C*) Solyc08g007530, *AHL1*. (*D*) Solyc03g098280, *SlAGO1b*, *ARGONAUTE 1b*. Panels in each figure part follow the descriptions for Figure 2.

DE sRNAs mapped to genes arranged in clusters across the tomato genome (figure 4, supplementary tables S4, S7, Supplementary Material online); this led to an increased number of genes per gene class that were consistently targeted by sRNAs. Therefore, the identity of genes that were observed in physical clusters drove our enrichment analyses. These physically linked gene families have undergone expansions in the *Solanum* lineage compared to *Arabidopsis* (supplementary tables S6, S7, Supplementary Material online). An example of this pattern are genes belonging to the protein Panther subfamily NUCLEAR TRANSPORT FACTOR 2/RNA RECOGNITION MOTIF PROTEIN (PTHR31413:SF10) with a single member in *Arabidopsis* (Mi et al. 2017). Thirteen of these genes were targeted mostly by 21-22-nt DE sRNAs, nine of them arranged in clusters on chromosome 2 (supplementary tables S4, S7, Supplementary Material online). Genes belonging to the Panther subfamily PROTEIN FLOWERING LOCUS D (PTHR10742:SF260) have likewise expanded in *Solanum*, with 14 members in contrast to the single copy in *Arabidopsis* (Mi et al. 2017). We found DE sRNAs mapping to 11 of these genes arranged in clusters on chromosome 5, some of which showed reduced expression levels in hybrid endosperm (supplementary tables S3, S4, S7, Supplementary Material online).

**FIG. 4.**
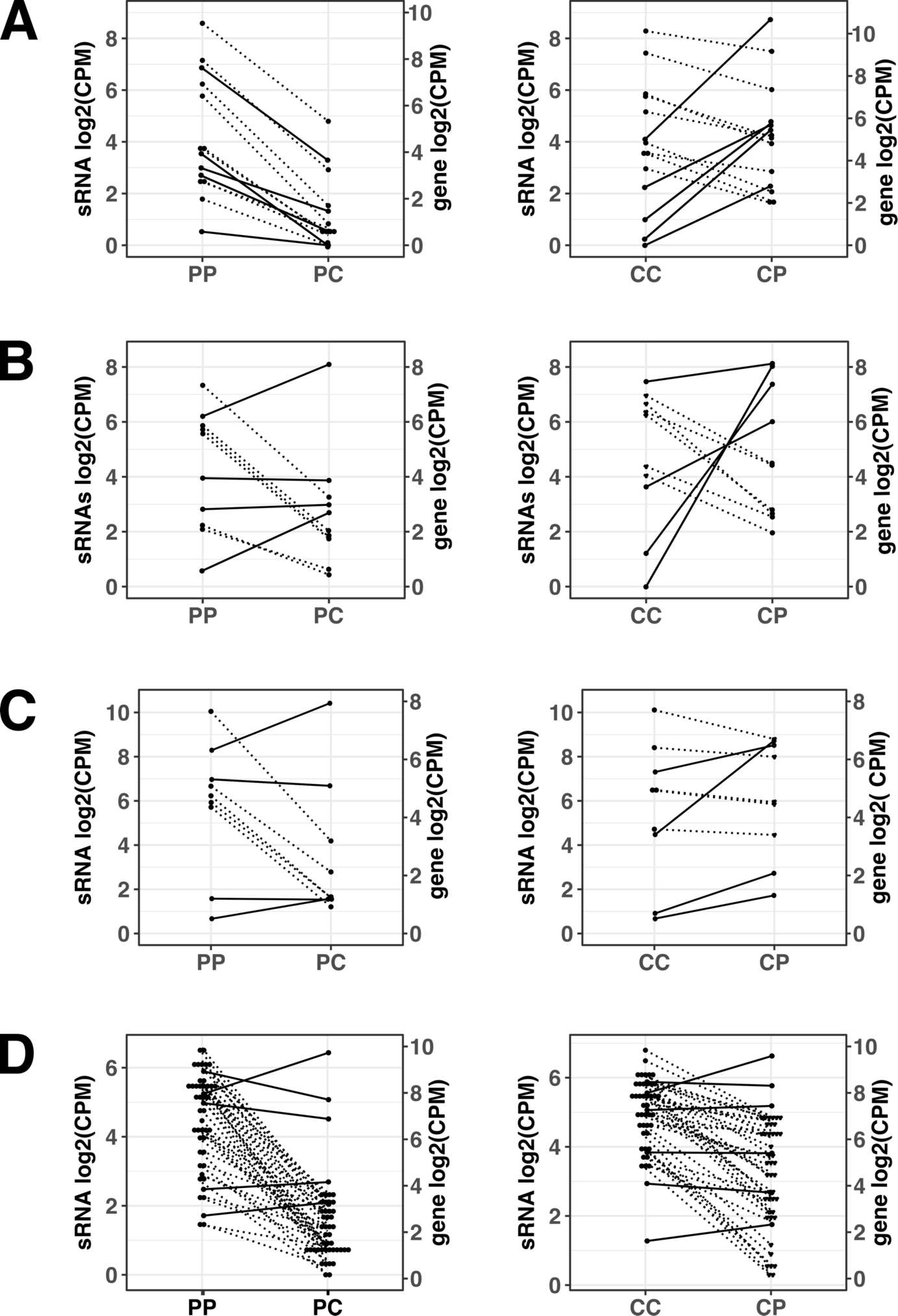
Examples of DE sRNA target genes arranged in clusters. (*A*) *ARID5* family Solyc12g038750, Solyc12g038740, Solyc12g094730, Solyc12g095740, Solyc01g017030, Solyc01g017790. (*B*) *CLASSY-3* family Solyc08g077690, Solyc01g068320, Solyc01g068300, Solyc01g060460. (*C*) Kinase family protein (D7MB90_ARALY) Solyc12g009370, Solyc12g016120, Solyc12g015990, Solyc05g007140. (D) YIPPEE domain (PTHR13848:SF5) gene family Solyc03g096110, Solyc03g096130, Solyc03g096120, Solyc03g096160, Solyc03g096140, Solyc03g096100, Solyc03g096260, Solyc03g096150, Solyc03g096250, Solyc03g096080. Panels in each figure part follow the descriptions for Figure 2.

Other genes arranged in clusters with sRNAs mapping to them are the chromatin remodelling protein families AT-RICH INTERACTIVE DOMAIN-CONTAINING PROTEIN 5-RELATED (PTHR15348:SF15) (Baba et al. 2011; Chandler et al. 2013), hereafter called ARID5 family, and SNF2 DOMAIN-CONTAINING PROTEIN CLASSY 3-RELATED, hereafter called CLASSY3 family (figure 4A, B; supplementary table S4, Supplementary Material online). Members of the CLASSY family have putative roles in RdDM (Law et al. 2011) and have recently been shown to be important regulators of sRNA production in *Arabidopsis* (Zhou et al. 2018). For many of these *ARID5* and *CLASSY3* family genes, the reduction in sRNA expression in the seed is invariably concurrent with markedly increased gene expression in the CP hybrid endosperm (figure 4A, B, right panels). This is not the case in PC hybrid endosperm, in which some genes belonging to the *ARID5* family exhibit reduced expression (figure 4A). Other members of gene families occurring in clusters and targeted by DE sRNAs include members of the Kinase family protein (D7MB90_ARALY) clustered on chromosome 12 (figure 4C), and genes with a YIPPEE domain (PTHR13848:SF5) clustered on chromosome 3 (figure 4D). The latter class of genes has been shown to play a role in epigenetic regulation of chromatin, with conditional knockout mouse lines resulting in hypomethylated DNA and embryonic lethality (Kim et al. 2012; Subramanian et al. 2016).

Some of the genes targeted by DE sRNAs did not exhibit any detectable expression. Lack of expression may indicate that these sRNAs inhibit transcription of these genes, possibly via RdDM or related mechanisms leading to TGS or PTGS (Matzke and Mosher 2014; Pikaard and Mittelsten Scheid 2014; Cuerda-Gil and Slotkin 2016). For example, only six of the ten genes encoding a YIPPEE domain targeted by DE sRNAs were expressed in endosperm (figure 4D, gene expression). Another example of clustered genes producing DE sRNAs but without detectable gene expression in endosperm is a cluster of *DICERLIKE* genes located on chromosome 1 (supplementary tables S4, S7, Supplementary Material online). DICER proteins are pivotal components of gene silencing pathways and involved in the cleavage of double-stranded RNA, thus producing siRNAs (Matzke and Mosher 2014; Pikaard and Mittelsten Scheid 2014; Borges and Martienssen 2015; Cuerda-Gil and Slotkin 2016). From the 48 *DICERLIKE* genes annotated in the SL2.50 version of the tomato genome, 39 are located in a 5.5-Mb stretch of chromosome 1, and six of these genes are targeted by significantly DE sRNA clusters in hybrid seeds (supplementary tables S4, S7, Supplementary Material online). Irrespective of the sRNA levels in seeds, there was no detectable endosperm expression of any of these DICER-coding genes clustered on chromosome 1. A single *DICER-LIKE 2c* gene, Solyc11g008520 located on chromosome 11, was consistently less expressed in hybrid endosperms of both cross directions (supplementary table S3, Supplementary Material online). Another set of genes with mapped DE sRNAs, yet without detectable gene expression (indicating possible sRNA-mediated TGS or PTGS), encodes proteins belonging to the family AMINOTRANSFERASE-LIKE, MOBILE DOMAIN PROTEIN-RELATED (PTHR46033:SF1); 10 genes were found on chromosome 4 with DE sRNA clusters mapping to them (supplementary tables S4, S7, Supplementary Material online). Genes of the same family in *Arabidopsis*, *MAIL1* and *MAIN*, have recently been shown to be involved in a gene silencing pathway independent of sRNAs and DNA methylation (Ikeda et al. 2017).

### Mode of Expression in Hybrids: Dominance May Reflect Differences in Levels of Parental Conflict

We assessed the mode of expression (conserved, additive, dominant, overdominant, or underdominant) of sRNAs and gene transcripts by comparing total expression levels in *S. peruvianum*, *S. chilense*, and their reciprocal hybrids. Following the rationale described in previous studies (McManus et al. 2010; Combes et al. 2015), we performed analyses of expression modes for the DE transcripts and sRNAs as well as for the whole set of transcripts and identified sRNA clusters (figure 5; supplementary figures S3, S4, Supplementary Material online). The analysis of expression modes of all expressed and genes and sRNA clusters (figure 5A, B) revealed that a large proportion of these show conservation of parental (within-species) expression levels in the hybrids, particularly for gene expression (>60%; figure 5A, purple). While conserved sRNA expression was also the dominant expression mode when evaluating all sRNAs (58.6% in *S. chilense* and 56.9% in *S. peruvianum*; figure 5B), the entire sRNA data set also revealed a marked pattern of non-conservedness, with maternal dominance being a major category (30.5% in *S. chilense* and 28.6% in *S. peruvianum*; figure 5B).

**FIG. 5.**
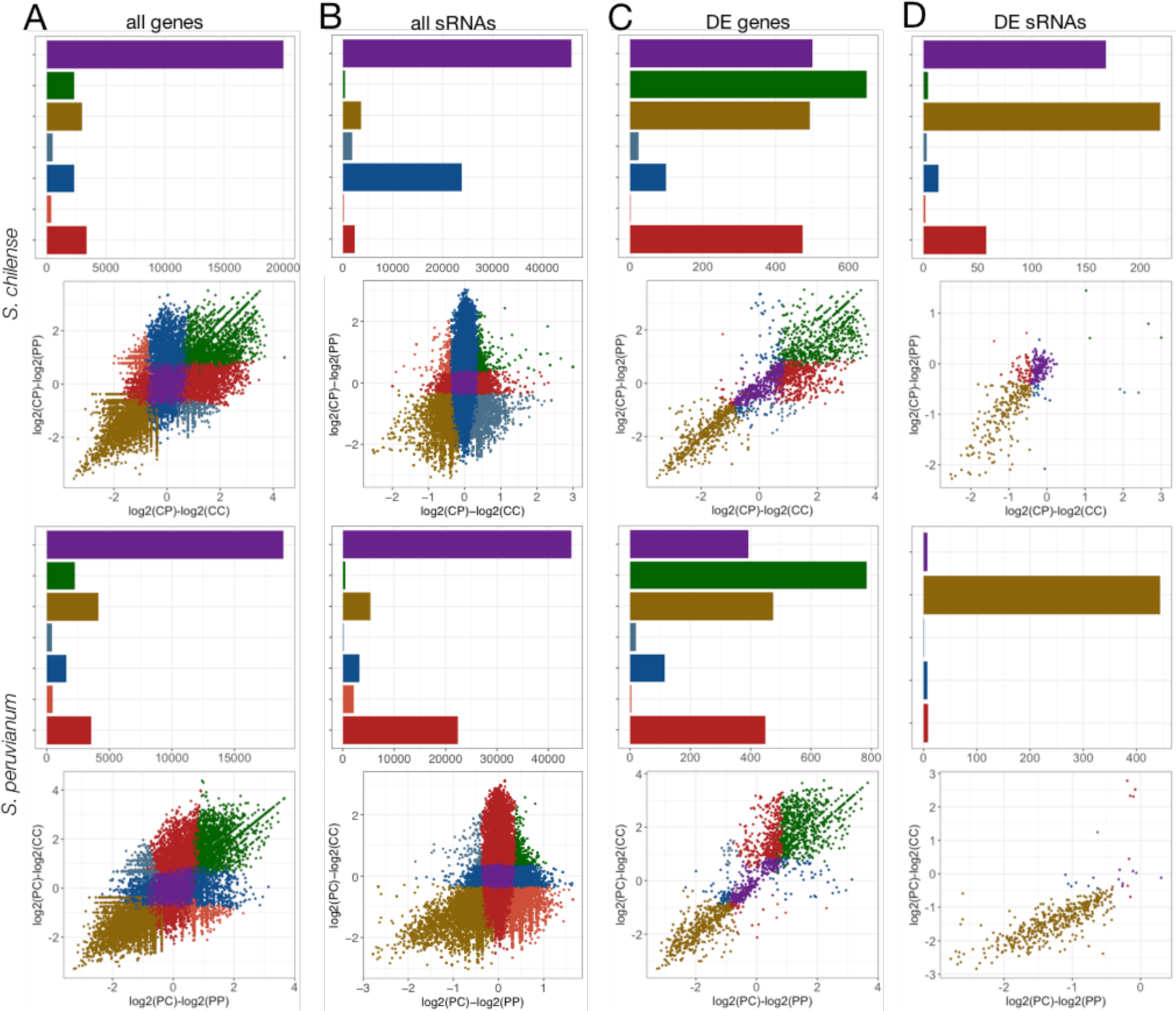
General patterns of expression modes. Hybrid *vs*. normal seed comparisons with *S. chilense* (upper panel) and *S. peruvianum* (lower panel) as maternal plants, respectively. (*A*) All genes (*n* = 33,805). (*B*) All sRNA clusters (*n* = 57,229). (*C*) Differentially expressed genes (*n* = 2,295). (*D*) Differentially expressed sRNA clusters (*n* = 468). Expression mode categories are colored as follows: conserved, purple; overdominant, green; underdominant, yellow; *S. chilense* additive, light blue; *S.chilense* dominant, dark blue; *S. peruvianum* additive, light red; *S. peruvianum* dominant, dark red.

We separately examined the expression modes of 21-22-nt sRNAs, 24-nt sRNAs, and miRNAs for the whole set of sRNAs as well as for DE sRNA clusters (supplementary figures S3, S4, Supplementary Material online). Most of the identified sRNA clusters were 24 nt (supplementary table S2; supplementary figure S3, Supplementary Material online). The levels of non-conserved expression modes were larger in the 21-22-nt category, with maternal dominance reaching 40.5% in *S. peruvianum* and 34.7% in *S. chilense*. The underdominant class was also larger in the 21-22-nt sRNA category than in the other two categories (supplementary figure S3*B*, Supplementary Material online).

Some DE genes in both species show transgressive expression (figure 5C), with overdominance being the predominant trend followed by underdominance of gene expression. Maternal dominance also markedly contributes to gene expression in the hybrids. An interesting result is the high proportion of genes that are in the *S. peruvianum*-dominant category in CP hybrid seeds, surpassing the maternal-dominant category for *S. chilense* (15.9% vs. 3.8%; figure 5C, upper panel). This result suggests that *S. peruvianum* in the paternal role greatly influences gene expression in CP hybrid endosperm despite contributing only one haploid genome. The signature of *S. peruvianum* dominance of gene expression in the CP hybrid is also evident in the expression mode of all genes and not only the DE genes (figure 5A, upper panel). Although most genes in both cross directions show a conserved pattern of expression, the *S. peruvianum* dominant category ranks second, surpassing other expression modes (10.5% *S. peruvianum* dominance; figure 5A, upper panel). These results indicate that the ‘genomic dominance’ of *S. peruvianum* relative to *S. chilense* is not restricted to DE genes but acts at a genome-wide level.

DE sRNAs were almost completely underdominant in PC hybrid seeds (figure 5D, supplementary figure S4, Supplementary Material online). In contrast, many DE sRNA clusters in CP hybrid seeds showed conserved expression. The *S. peruvianum*-dominant signature evident in the mode of gene expression (figure 5A, C) is also apparent in the expression mode of DE sRNA clusters in CP hybrid seeds, with 11.6% of the DE sRNA clusters falling into this category. However, the most striking trend in the expression mode of sRNAs was that of underdominance of DE sRNA clusters in hybrid seeds (figure 5D, supplementary figure S4, Supplementary Material online).

## Discussion

### Evidence for Conserved Epigenetic Landscapes in Compromised Hybrid Endosperm

Our analyses of sRNAs and transcripts that were differentially expressed between normal and failing seeds/endosperms revealed striking similarities with previous work on transcriptomic responses to hybridization in other taxa, particularly with the effects of Pol IV mutations on the epigenomic landscape of *Arabidopsis* endosperm. Erdmann et al. (2017) demonstrated that the Pol IV sRNA pathway mediates dosage interactions between maternal and paternal genomes. Specifically, they showed that disabling mutations in *nrpd1* led to shifts toward higher expression of maternally inherited alleles. These results mirror our previous findings of increased maternal expression proportions in *Solanum* hybrid endosperms (Florez-Rueda, Paris, et al. 2016). Likewise, Erdmann et al. (2017) reported increased gene expression in *nrpd1* mutant endosperm compared to WT, resembling the increased gene expression among DE genes in *Solanum* hybrid endosperms (figure 1A).

Taken together, the reduction in Pol lV expression and the overall increase in expression of DE transcripts and maternal expression proportions in hybrids (Florez-Rueda, Paris, et al. 2016) allows us to draw comparisons between effects of the *Arabidopsis nrpd1* mutant (Erdmann et al. 2017) and the natural case of HSF we explore here in *Solanum*. Based on these obvious parallels, we postulate a *Solanum* Pol lV sRNA pathway acting in a similar fashion to that described in *Arabidopsis* (Erdmann et al. 2017), mediating dosage interactions of the parental genomes upon fertilization. We propose that the Pol lV sRNA pathway serves to maintain the expected 2:1 ratio in the endosperm, likely through direct and/or indirect effects on many genes in the endosperm. The observed reduced expression of the main Pol IV subunits may be functionally linked to the increased maternal expression proportions in hybrid endosperm of wild tomatoes (Florez-Rueda, Paris, et al. 2016).

Increased expression of MADS-box TF genes upon hybridization has previously been reported in *Arabidopsis* (Josefsson et al. 2006; Walia et al. 2009; Hehenberger et al. 2012; Lu et al. 2012; Burkart-Waco et al. 2013), *Capsella* (Rebernig et al. 2015), and *Oryza* (Ishikawa et al. 2011). We found a large number of MADS-box genes (among other TF genes) overexpressed in both reciprocal hybrid endosperms (supplementary figure S2*E*, supplementary table S3, Supplementary Material online). MADS-domain TFs have been shown to play key regulatory roles in plant reproduction, in particular in regulating female gametophyte, embryo, and endosperm development (reviewed in Masiero et al. 2011). Likewise, *AGAMOUS-LIKE* (*AGL*) genes were jointly overexpressed in ‘paternal-excess-like’ crosses involving *S. chilense*, *S. peruvianum*, and *S. arcanum* (Roth et al. 2019). These TFs are part of the GO protein dimerization activity (GO:0046983) and include 11 *AGL* genes, 13 *2FE-2S FERREDOXIN-LIKE* genes, three *PHERES* genes, *APETALLA3* and *SEPALATA3*, among others (supplementary tables S3, S6, Supplementary Material online). AGL proteins, which themselves are MADS-domain proteins, have been shown to affect endosperm development in *Arabidopsis* (Kang et al. 2008; Shirzadi et al. 2011). Intriguingly, overexpression of *AGL62* and *AGL90* is associated with the postzygotic barrier between *A. thaliana* and *A. arenosa*, which manifests as endosperm over-proliferation and delayed development (Josefsson et al. 2006; Walia et al. 2009; Burkart-Waco et al. 2013). Providing functional validation for this pattern, transgenic underexpression of *AGL62* attenuated the level of HSF in *Arabidopsis* (Hehenberger et al. 2012).

sRNAs have been shown to modulate expression of MADS-box TF genes; maternal small interfering RNA (siRNA) expression is negatively correlated with *AGL* gene expression in *Arabidopsis* endosperm (Lu et al. 2012). However, our analyses did not support a consistent trend of sRNAs targeting MADS-box TF genes; we found but a single gene Solyc12g056460 (*SOC1*) with an associated DE sRNA cluster (supplementary table S4, Supplementary Material online). Taken together, this and earlier *Arabidopsis* studies suggest that the putative functions of MADS-domain TFs in mediating both normal seed development and endosperm-based HSF are conserved across angiosperms. Specific functions of MADS-box TF genes in *Solanum* have not yet been studied, but here we have uncovered a list of candidates with potentially important roles that remain to be functionally validated.

Qualitative and quantitative sRNA differences between the parental genomes may affect the expression of genes and TEs neighboring the sRNAs in hybrid seeds. In some instances of hybridization, changes in sRNA expression are concomitant with heterosis (Groszmann et al. 2011; Barber et al. 2012), although a causal role of sRNAs has not been shown; in our *Solanum* case and others, they may lead to HSF (Ng et al. 2012; Kirkbride et al. 2015; Florez-Rueda, Paris, et al. 2016; Garner et al. 2016). We found that DE sRNAs were consistently underexpressed in hybrid seeds (figure 1B); this trend is reflected in underdominance of sRNA expression in hybrid seeds when compared to seeds derived from intraspecific crosses on the same maternal plant. Underdominance of sRNA expression upon hybridization has been reported in other tissues besides the seed and in hybridization cases in diverse plant genera (Groszmann et al. 2011; Barber et al. 2012; Lu et al. 2012; Shen et al. 2012; Shivaprasad et al. 2012; He et al. 2013). In all these cases, as well as ours, the molecular mechanisms leading to reduced sRNA levels are still unknown; based on the reduced expression of Pol IV subunits (supplementary table S4, Supplementary Material online), we can hypothesize that perturbations in the Pol lV sRNA pathway may be involved (Erdmann et al. 2017).

We uncovered high levels of maternal dominance of sRNA expression that may be explained by the nature of the seed tissue we collected (seeds extracted manually with subsequent washes), maternal seed coat being one of its components. Another possible scenario is that the sRNAs exhibiting maternal dominance may be generated by filial seed tissues. However, there is disagreement among studies in *A. thaliana* whether 24-nt siRNAs show strongly maternally-biased expression (Mosher et al. 2009; Erdmann et al. 2017; Kirkbride et al. 2019). Regardless, these sRNAs are thought to accumulate in the endosperm and to mediate gene expression (Calarco and Martienssen 2011); the high level of observed maternal dominance in the expression inheritance of sRNAs in both species suggests that this may also be the case in *Solanum*.

### Feedback Regulation of Core Silencing Proteins through sRNA-mediated Silencing

Our data suggest that sRNAs that are DE in hybrid seeds target many genes with important functions in sRNA biogenesis and epigenetic regulation. Importantly, members of the *ARID5* and *CLASSY3* families, *DICER*, *AGO1B*, *AGO5*, *RDM1*, and *DMS3* were shown to be associated with sRNAs in tomato seeds in abundances that were significantly different in both PC and CP hybrid seeds (figures 2–4, supplementary table S4, Supplementary Material online). For some of these genes, we were able to additionally assess gene expression levels; the apparent effect of sRNA abundance on gene expression suggests that sRNA-mediated gene silencing impacts the expression of some of these genes and may be defective in hybrid seeds, plausibly contributing to HSF. We hypothesize that these genes, some of which are regulators of TGS or PTGS themselves, are subject to feedback regulation orchestrated by their own sRNA products. Negative feedback regulation of *DICERLIKE* genes has been described in *Arabidopsis* (Xie et al. 2003; Bologna and Voinnet 2014) and yeast (Oberti et al. 2015); such feedback regulation is thought to allow homeostatic control of the cellular silencing machinery (Bologna and Voinnet 2014). The only gene for which we detected an effect on allele-specific expression is the ARGONAUTE-encoding gene Solyc03g098280, *SlAGO1b* (figure 3D). As a paternally expressed gene (PEG) with a low maternal proportions in normal endosperm of *S. peruvianum*, this gene showed the ‘typical’ increase (from 0.25 to 0.87 maternal proportion) in PC hybrid endosperm that we previously uncovered for the majority of PEGs (Florez-Rueda, Paris, et al. 2016). We posit that the observed underexpression of sRNA clusters mapping to *SlAGO1b* and its flanking regions may be responsible for its increased gene expression, with a higher maternal proportion in hybrid endosperm derived from maternal *S. peruvianum*.

Although we cannot provide functional verification to support feedback regulation of genes involved in sRNA-mediated gene silencing in *Solanum* endosperm, our results provide pioneering glimpses into the epigenetic landscape in the context of HSF. We could show that DE sRNA clusters map to genes playing pivotal roles in epigenetic regulation, with expected implications for HSF. Further characterization of the epigenomic landscape of the endosperm through chromatin immunoprecipitation and sequencing (ChIP-seq) as well as methylome sequencing will allow a proper evaluation of these hypotheses.

### Mode of Expression in Hybrids: Dominance may reflect Differences in Parental Conflict

Previous analyses of expression modes have been restricted to evaluating inheritance in whole plants that were successful hybridization products of within- or among-species crosses (Eichten et al. 2011; Shi et al. 2012; Bell et al. 2013; Combes et al. 2015; Li et al. 2015; Carlson et al. 2017). Although these types of analyses on whole hybrid plants provide valuable insights into the transcriptomic effects of hybridization, they do not address the issue of parental conflict that is expected to play out in the developing seed (Haig and Westoby 1991; Haig 2013; Lafon-Placette and Köhler 2016).

The near-complete HSF phenotype characterizing both cross directions between *S. peruvianum* and *S. chilense* (yet with marked phenotypic differences between reciprocal crosses) may be seen as resulting from different levels of parental conflict among diverged parental lineages (Haig and Westoby 1989; Brandvain and Haig 2005; Haig 2013). Hybrid seeds from *S. chilense* maternal plants (CP) are larger, showing a ‘paternal excess-like’ phenotype in contrast to the smaller hybrid seeds with *S. peruvianum* mothers (PC) that show a ‘maternal excess-like’ phenotype (Florez-Rueda 2014; Florez-Rueda, Paris, et al. 2016; Roth, Florez-Rueda, Griesser, et al. 2018). The reciprocal differences both in early seed development and mature hybrid seed size suggest that the *S. peruvianum* lineage has evolved under higher levels of parental conflict than has *S. chilense.* These patterns and inferences are consistent with higher range-wide nucleotide diversity, indicative of higher effective population size (Städler et al. 2008; Tellier et al. 2011; Beddows et al. 2017), and higher expression levels of imprinted genes in *S. peruvianum* (Roth, Florez-Rueda, Paris, et al. 2018).

Likewise, *S. peruvianum* drives expression landscape polarization in hybrid endosperms derived from reciprocal crosses with *S. chilense* and *S. arcanum* (Roth et al. 2019). In line with these observations, our analyses of the expression modes of DE sRNAs and genes revealed a trend of *S. peruvianum* dominance in CP hybrid seeds and endosperm, respectively (figure 5C, D, upper panel). This signature held true not only for DE genes and sRNAs but was also detected at a genome wide-level, specifically on the larger dataset of all expressed genes where the *S. peruvianum*-dominant category ranked second (figure 5A, upper panel). We interpret the pattern of *S. peruvianum* dominance as consistent with the rationale of the weak inbreeder/strong outbreeder (WISO) hypothesis (Brandvain and Haig 2005), with the *S. peruvianum* genome ‘overpowering’ that of *S. chilense*, which putatively evolved under lower levels of parental conflict. These inferences are in accordance with our prior and current evidence for higher effective ploidy of *S. peruvianum* compared to *S. chilense* (Roth et al. 2019), and how it plausibly underpins seeds’ developmental and phenotypic differences between these two wild tomato lineages: smaller hybrid seed size and transcriptomic signals (Florez-Rueda, Paris, et al. 2016; Roth, Florez-Rueda, Griesser, et al. 2018; Roth et al. 2019), such as a higher proportion of DE genes, DE sRNAs, and larger shifts toward maternal allelic expression in PC (compared to CP) hybrid endosperm.

### Data Availability

Raw sequence data for the RNA-sequencing dataset used in this study are available from the Sequence Read Archive (https://trace.ncbi.nlm.nih.gov/Traces/sra/) with the accession numbers XXXXXXXXX (sRNAs; this study) and SRX1850236 (mRNA; Florez-Rueda, Paris, et al. 2016).

## Acknowledgments

We are grateful to Maja Frei and Esther Zürcher for taking expert care of the plants, to Margot Paris and Anja Schmidt for technical advice, and to Alex Widmer for general support of this project. We thank the C.M. Rick Tomato Genetics Resource Center at U.C. Davis for generously supplying seed samples, and Lennart Opitz for bioinformatics support. We also thank the Genetic Diversity Center (ETH Zurich, Switzerland) and the Swiss Institute for Bioinformatics (Lausanne, Switzerland) for providing valuable tools and training for bioinformatics analyses. This work was supported by the University of Zurich, the ETH Zurich, and grants from the Swiss National Science Foundation [grant number 31003A_130702 to T.S., 310030B_160336 to U.G.], an ETH Research Grant [grant number ETH-40 13-2 to T.S. and Alex Widmer], and the University of Zurich Research Priority Program Evolution in Action [Pilot Grant to A.M.F.-R. and U.G.].

