## Supplementary Figures for "Endosperm and seed transcriptomes reveal possible roles for small RNA pathways in wild tomato hybrid seed failure"

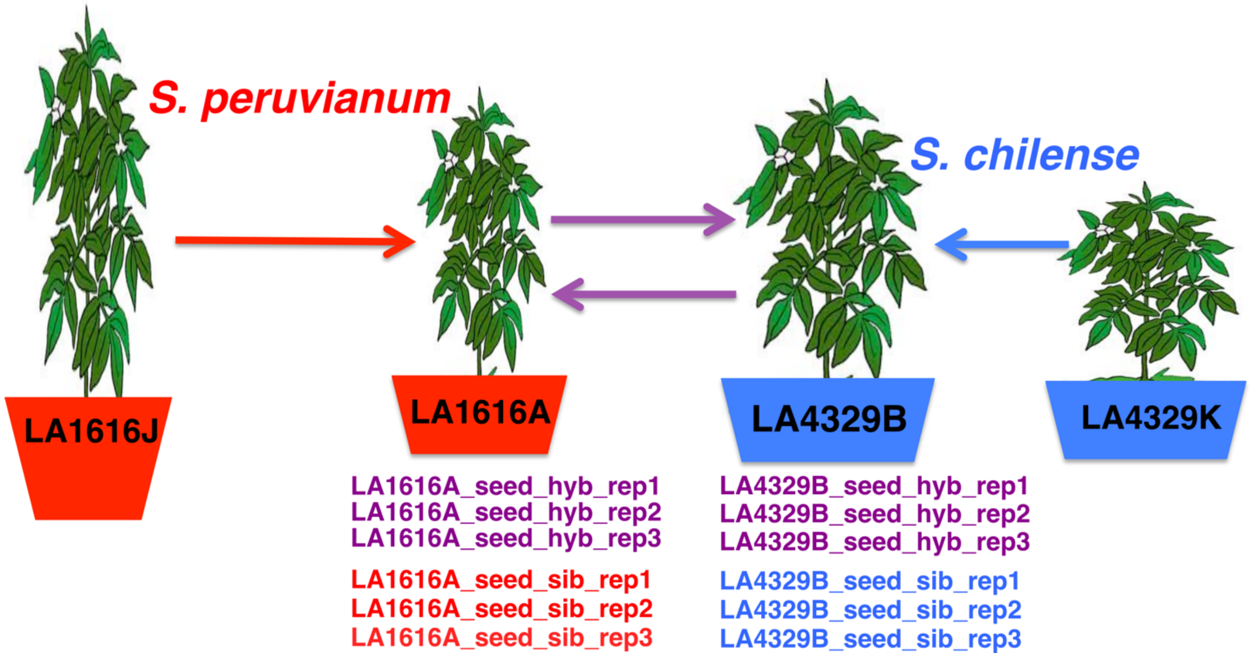


**FIGURE. S1**—Small-RNA crossing design and libraries used. For each of the main plants, namely LA1616A and LA4329B, inter- and intra-specific crosses were performed. Plants LA1616J and LA4329K were only used as pollen donors in intraspecific crosses ('sib' libraries). Small-RNA libraries were prepared in triplicate per cross type and cross direction. Arrows indicate the direction of manual pollinations.

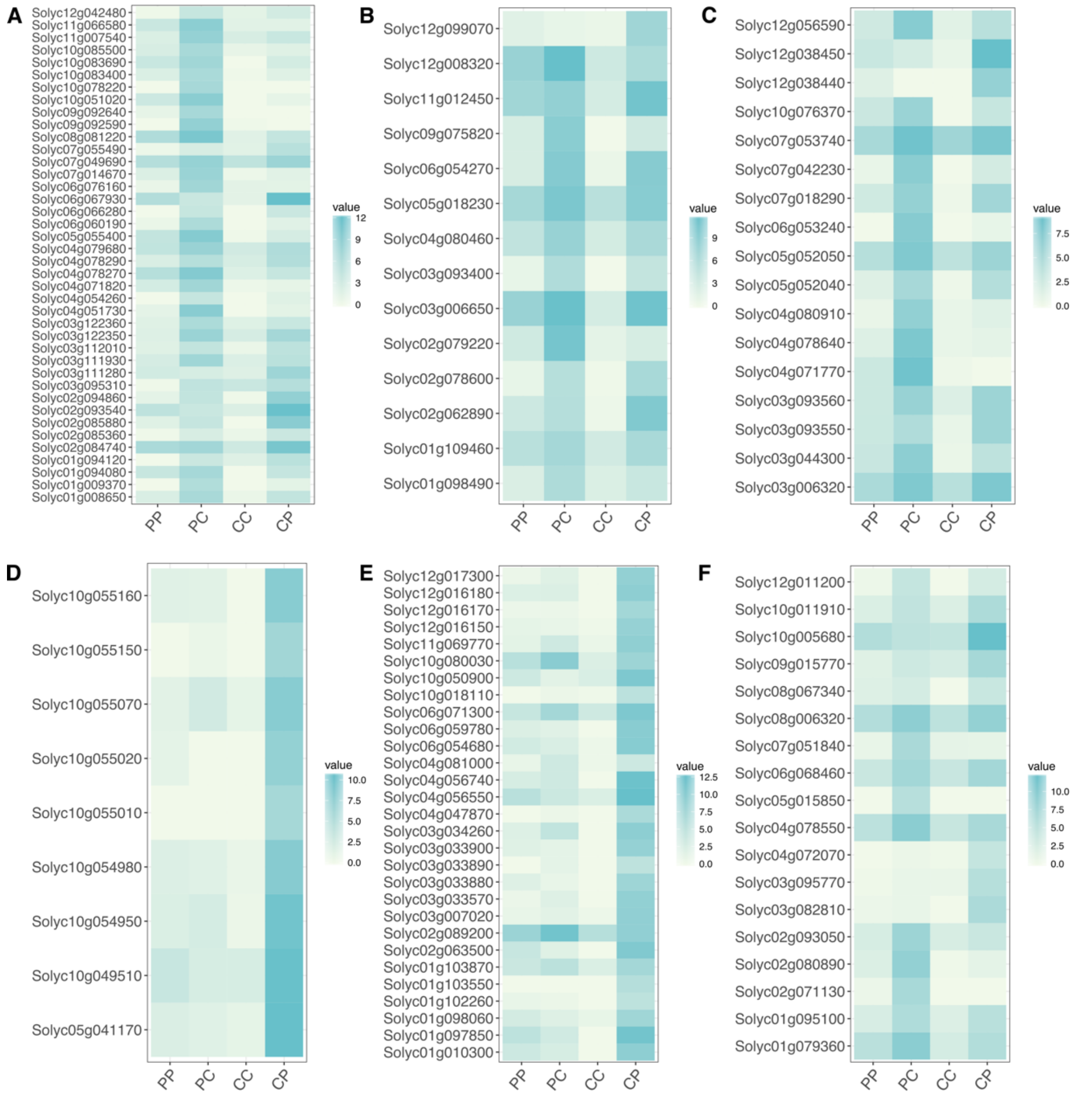


**FIGURE. S2**—Enriched pathways upregulated in hybrid endosperm. Heatmaps of significantly differentially expressed genes in the following domain families: (*A*) PF00067-cytochrome P450; (*B*) PF00083-sugar transporter; (*C*) IPR001471-AP2/ERF domain; (*D*) IPR013921-mediator complex; (*E*) IPR002100-transcription factor, MADS-domain; (*F*) IPR003657-WRKY domain.

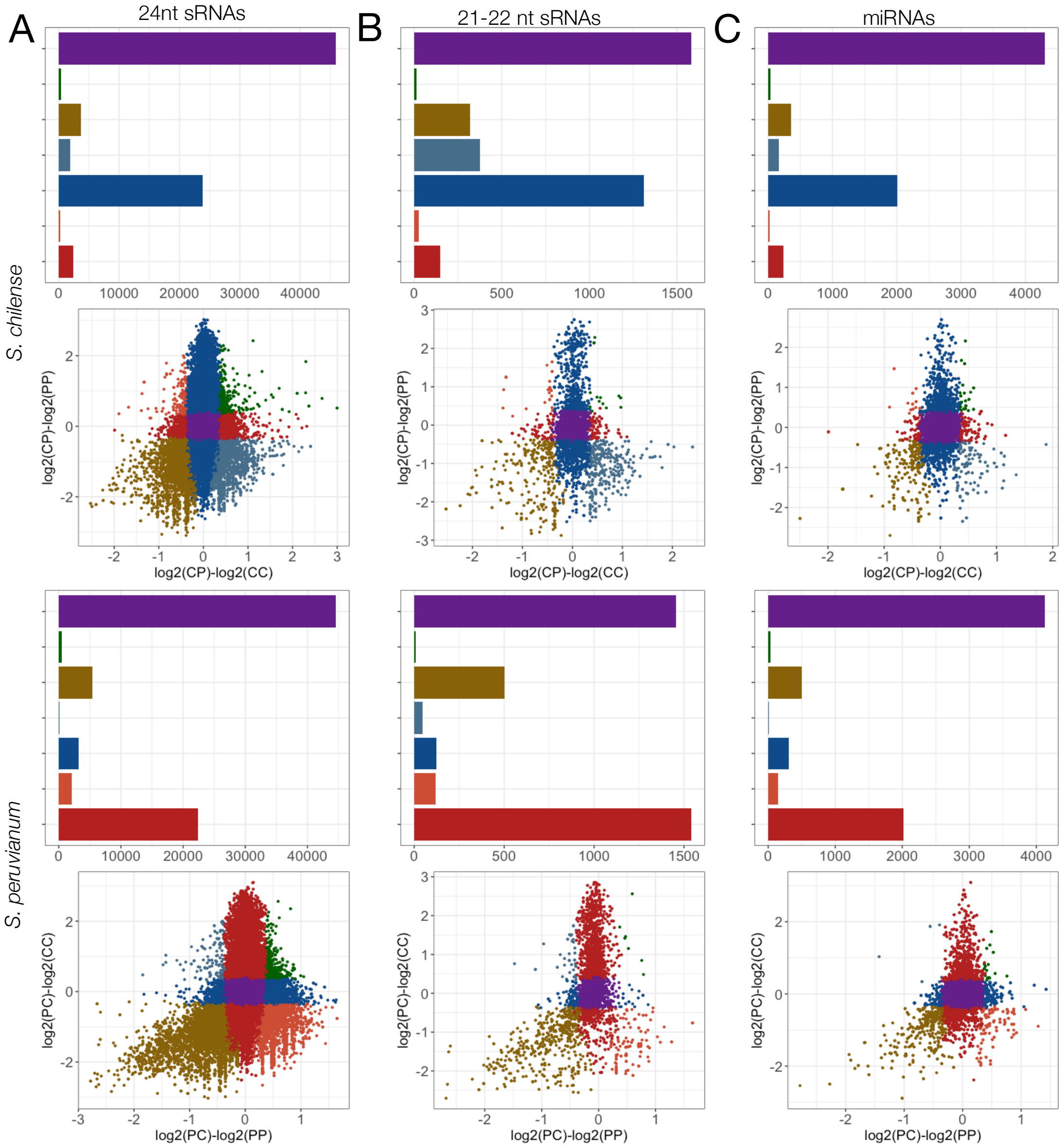


**FIGURE. S3**—Inferred expression modes for small-RNA cluster categories. Hybrid *vs.* normal seed comparisons with *S. chilense* (upper panel) and *S. peruvianum* (lower panel) as maternal plants, respectively. (*A*) 24 nt sRNAs ($n$ = 51297); (*B*) 21-22 nt sRNAs ($n$ = 2958); (*C*) miRNAs ($n$ = 2974). Color coding of expression mode categories is the same as for figure 5.

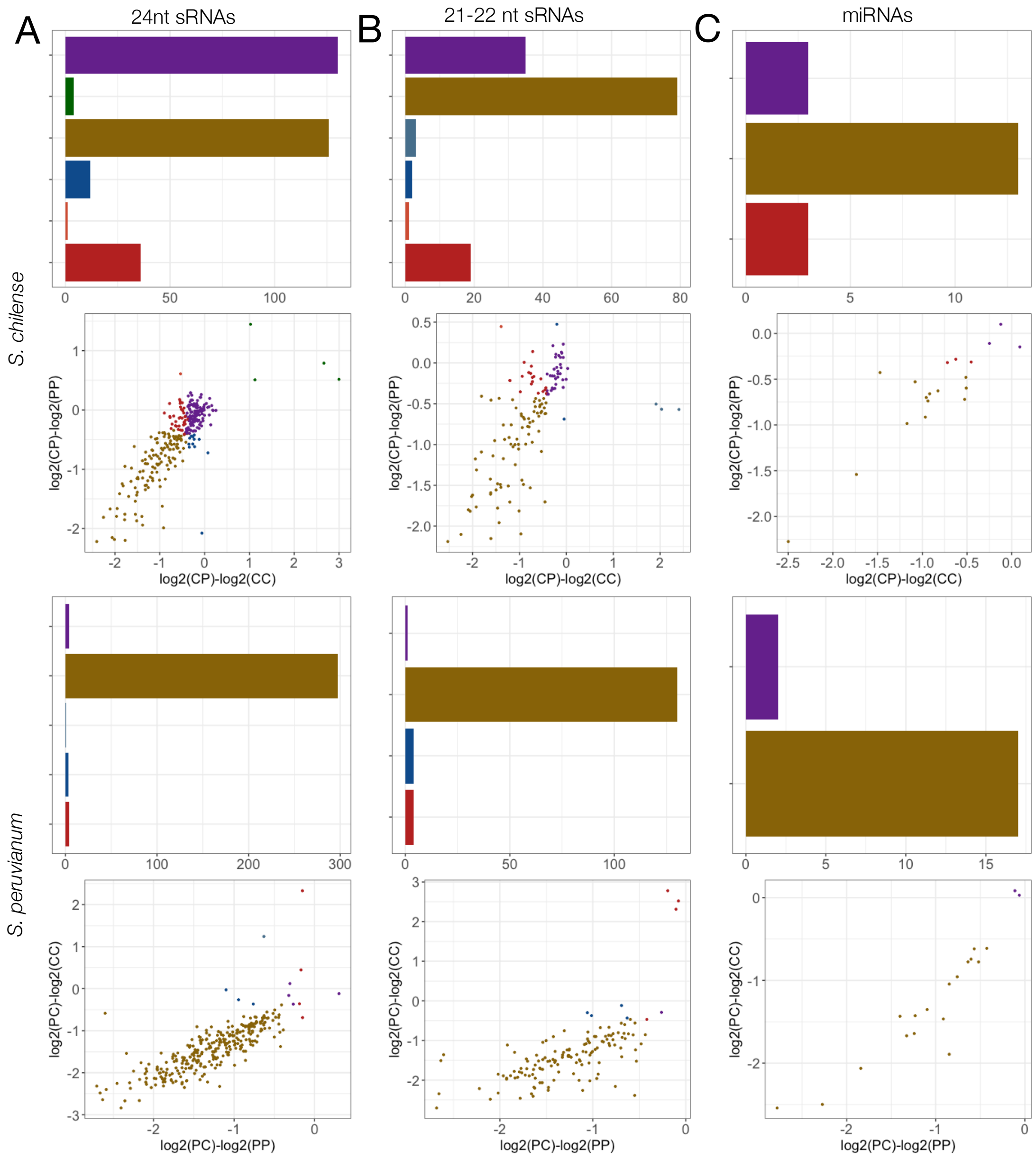


**FIGURE. S4**—Inferred expression modes for differentially expressed sRNAs. Hybrid *vs*. normal seed comparisons with *S. chilense* (upper panel) and *S. peruvianum* (lower panel) as maternal plants, respectively. (*A*) 24 nt sRNAs ($n$ = 310); (*B*) 21-22 nt sRNAs ($n$ = 139); (*C*) miRNAs ($n$ = 19). Color coding of expression mode categories is the same as for figure 5.
